# Detection of protein markers for autophagy, autolysis and apoptosis processes in a *Saccharomyces cerevisiae* wine flor yeast strain when forming biofilm

**DOI:** 10.1101/324772

**Authors:** Jaime Moreno-García, Juan Carlos Mauricio, Juan Moreno, Teresa García-Martínez

## Abstract

Yeast autophagy, autolysis and apoptosis are biological processes triggered by nutrient starvation conditions that usually take place in winemaking. Biological aging of Sherry wines constitutes an enological environment suitable for the induction of these mechanisms due to the scarcity of nutrients and formation of yeast social communities, i.e.; biofilm; however, few studies have been carried out in this regard. Here, we perform a proteomic analysis to detect any autolysis/autophagy/apoptosis protein markers and/or proteins potentially related to these processes under flor forming and fermentative conditions. The scarce presence of autophagy proteins in flor biofilm forming conditions, the existence of autophagy inhibitors (e.g. Pph21p), and high quantity of crucial proteins for autolysis and apoptosis, Pep4p and Mca1p, respectively; indicate that autophagy may be silenced while autolysis and apoptosis are activated when the yeasts are forming flor. This is the first time that autophagy, autolysis and apoptosis proteomes have been studied as a whole in flor yeast under film forming conditions to our knowledge.

## 1. Introduction

Yeast autolysis has been a subject of study for decades and its positive influence on the organoleptic profile of some types of wines is widely recognized (reviewed in [1,2]), while biological processes of autophagy and apoptosis are less known in wine yeasts (reviewed in [3–6]). These three processes —autophagy, autolysis and apoptosis— are associated with each other and are all triggered by starvation for nutrients (among other stress conditions) [7–12], which usually happen in enological environments. Autolysis, for instance, takes place during ageing of sparkling wines produced by the traditional méthode champenoise when yeasts are subjected to low contents of nitrogen and carbon conditions [2,13]. Autophagy was reported as well during sparkling wine aging by the occurrence of morphological changes and the presence of autophagic bodies [14]. Piggot et al. (2011) [15] also showed that autophagy takes place in still wine fermentation. More recently, Orozco et al. (2013) [16] found that factors relating to apoptosis, such as caspase Mca1p or apoptosis-inducing factor Aif1p play a positive role in yeast longevity during winemaking in times of dwindling resources.

Other environments common in winemaking in which these processes may occur are flocs and biofilms. The benefit of a cellular suicide program in social communities seems evident because self-destruction of damaged and old cells, which consume dwindling nutrients, contributes to the viability/reproductive success of healthier members of the community. In the case of Sherry wine, special *Saccharomyce cerevisiae* strains, known as flor yeasts, have to deal with lack of fermentable carbon sources (among other stresses) and form an air-liquid biofilm formation, so-called flor, in biological aging conditions that occur after fermentation [17,18]. In this work, we attempt to approach the three biological processes —autophagy, autolysis and apoptosis— in a flor biofilm environment. Hitherto, only autolysis has been evidenced under biological aging conditions [19] while autophagy and apoptosis have remained unreported. Nonetheless, in a previous study, our group accounted for the presence of apoptosis factors when studying the mitochondrial proteome in a flor yeast when forming flor [20].

Following other authors’ experiments in which proteins are used as markers [21–24], we performed a targeted proteomic analysis to detect any autophagy/autolysis/apoptosis markers and/or related proteins, in a flor yeast strain under a biofilm forming condition (lacking glucose and high in ethanol) and under a fermentative condition (high glucose). This study is part of a sequence of un-targeted/targeted proteomic researches of flor yeasts [20,25–29], which distinctively analyze the autolysis/autophagy/apoptosis proteome under biofilm forming and fermentative conditions.

## 2. Materials and Methods

### Microorganism and cultivation conditions

*S. cerevisiae* G1 (ATCC: MYA-2451), a wild type of an industrial wine flor yeast strain, capable of fermenting and aging wine, from the Department of Microbiology (University of Cordoba, Spain) collection was used in this work. G1 under biological aging conditions, produces a thick flor velum about 30 days after inoculation with a cellular viability higher than 90% and a small proportion of sediment cells in the bottom of flasks [30].

A medium mimicking a biological aging condition, in this case a biofilm forming condition (BFC), was prepared without sugars consisting of 0.67% w/v YNB w/o amino acids, 10 mM glutamic acid, 1% w/v glycerol and 10% v/v ethanol, incubated at 21 °C without shaking for 29 days. Fermentative condition (FC) was developed in a medium containing 0.67% w/v YNB without amino acids, 10 mM glutamic acid, and 17% w/v glucose, and yeasts were incubated at 21 °C under gentle shaking for 12 h or until the middle of the log phase. 1×10^6^ 100 cells/mL were inoculated in each medium. All experiments were carried out by triplicate in flasks closed with hydrophobic cotton.

### Proteome analysis

Sampling times were chosen to obtain the maximum number of proteins in viable cells [30–33]. These were at the middle of the log phase, different for each condition: 12 hours from inoculation for FC and 29 days for BFC. At day 29th G1 flor yeast cells are in the initial phase of velum formation (Ph I) and the biofilm is completely formed in the air–liquid interface [34]. Methods for harvesting the cells and protein extraction are indicated in [20,25].

Yeast proteins under both conditions were extracted and later subjected to fractionation through 3100 OFFGEL (Agilent Technologies, Palo Alto, CA) followed by an identification by LTQ Orbitrap XL mass spectrometer (Thermo Fisher Scientific, San Jose, CA, USA) equipped with a nano LC Ultimate 3000 system (Dionex, Germany) (see [21–25] for more details). After identification, proteins were quantified in terms of the exponentially modified protein abundance index (emPAI; [35]).

Proteins related to autophagy, autolysis and apoptosis were selected by using SGD (http://www.yeastgenome.org/), Uniprot and references. These proteins together with the identification and quantification values are shown in Supplemental material 1. This file shows information about each autophagy, autolysis and apoptosis related protein detected in this analysis including a brief description, biological process and molecular function, the molar weight (Mr), a score value (combination of the XCorr values for its constituent peptides), observable and observed peptides and relative content as calculated from its PAI value. Protein content averages in mol% considering all proteins detected in each sample, were 0.24 at BFC and 0.16 at FC.

Information about proteins annotated in the autophagy, autolysis and apoptosis processes, considering the whole proteome of *S. cerevisiae* (https://www.yeastgenome.org/) and their content according to Ghaemmaghami et al. (2003) [31], were used as reference material (Table 1). Further, the SGD tool “GO Term finder” was used to determine the FDR (False Discovery Rate) and p-value for each protein group annotation considering all autophagy, autolysis and the apoptosis proteins in each sample (Supplemental material 2). p-value is defined at the probability or chance of seeing at least “x” number of genes (in our case ORFs) out of the total “n” genes in the list annotated to a particular GO (Gene Ontology) term, given the proportion of genes in the whole genome that are annotated to that GO Term. GO Terms with p-values lower than 0.1 have been highlighted (Supplemental material 2). The p-value is calculated using the Hypergeometric distribution. Four numbers are used to calculate each p-value: n, the number of objects in the sample; N, the number of objects in the reference population (6604 proteins from the *S. cerevisiae* whole proteome), k, the number of objects annotated with this item in the sample; and M, the number of objects annotated with the item in the reference population:

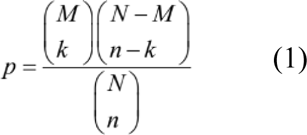

**TABLE 1.** Frequency and content of proteins related with autophagy, autolysis and apoptosis identified in flor yeast cells under biological forming (BFC) and under fermentative conditions (FC). Frequency of related proteins in *S. cerevisiae* whole proteome has been included. Protein content at log phase under rich-medium conditions [31] was included as reference material.

| Process |  | *S. cerevisiae* proteome [33] |  | FC |  | BFC |  |
| --- | --- | --- | --- | --- | --- | --- | --- |
|  |  | Protein frequency | Protein content (mol%) [29] | Protein frequency | Protein content (mol%) | Protein frequency | Protein content (mol%) |
| Total Proteins |  | 6721 |  | 611 |  | 413 |  |
| Autophagy | Total proteins | 95 (1.41%) | 0.83 | 11 (1.64%) | 1.25 | 6 (1.45%) | 0.40 |
|  | Regulation of induction: regulators | 26 (0.39%) | 0.20 | 4 (0.65%) | 0.18 | 2 (0.73%) | 0.17 |
|  | Regulation of induction: autophagosome-generating machinery | 24 (0.36%) | 0.12 | 1 (0.16%) | 0.09 | 0 (0%) | 0 |
|  | Cargo packaging | 8 (0.12%) | 0.32 | 2 (0.33%) | 0.57 | 2 (0.48%) | 0.04 |
|  | Vesicle nucleation | 5 (0.07%) | 0.01 | 0 (0%) | 0 | 0 (0%) | 0 |
|  | Vesicle expansion and completion | 26 (0.39%) | 0.15 | 1 (0.16%) | 0.32 | 1 (0.24%) | 0.19 |
|  | Retrieval | 7 (0.1%) | 0.03 | 1 (0.16%) | 0.01 | 1 (0.24%) | 0.01 |
|  | Docking and fusion | 14 (0.21%) | 0.04 | 2 (0.33%) | 0.07 | 0 (0%) | 0 |
|  | Vesicle breakdown | 1 (0.01%) | 0.01 | 0 (0%) | 0 | 0 (0%) | 0 |
|  | Permease efflux | 1 (0.01%) | 0 | 0 (0%) | 0 | 0 (0%) | 0 |
| Autolysis | Total proteins | 128 (1.90%) | 1.17 | 12 (1.96%) | 1.66 | 12 (2.91%) | 4.20 |
|  | Vacuolar proteases | 12 (0.18%) | 0.13 | 5 (0.82%) | 0.63 | 4 (0.97%) | 1.97 |
|  | Protease inhibitors | 4 (0.06%) | 0.003 | 0 (0%) | 0 | 2 (0.48%) | 0.32 |
|  | Glucanases | 12 (0.18%) | 0.19 | 2 (0.33%) | 0.67 | 2 (0.48%) | 1.48 |
|  | Nucleases | 108 (1.61%) | 0.65 | 5 (0.82%) | 0.36 | 2 (0.48%) | 0.07 |
|  | Mannosidases | 8 (0.12%) | 0.04 | 0 (0%) | 0 | 1 (0.24%) | 0.23 |
|  | Lipases | 32 (0.48%) | 0.16 | 1 (0.16%) | 0.01 | 1 (0.24%) | 0.12 |
| Apoptosis | Total proteins | 39 (0.58%) | 2.03 | 8 (1.31%) | 1.98 | 10 (2.42%) | 2.96 |

## 3. Results

Proteins involved in the autophagy, autolysis and apoptosis processes were identified in both BFC and FC samples. In both conditions, the frequency of proteins involved in these processes was high (> *S. cerevisiae* proteome frequency values, see Table 1). Apoptosis protein frequencies overpassed *S. cerevisiae* proteome frequency by 4 and 2-fold under BFC and FC, respectively. Autophagy proteins were more frequent and abundant under FC (considering the sum of content of all autophagy proteins under the conditions) (Table 1). The total protein content in BFC was half of the *S. cerevisiae* autophagy proteome content reported by Ghaemmaghami et al. (2003) [31] at log phase under rich-medium conditions pointing out a down-regulation during biological aging. The opposite happens for the autolysis and apoptosis proteomes in which BFC proteome was higher in frequency and abundance: 2.91 at BFC vs. 1.96% at FC and 4.20 at BFC vs. 1.66 mol% at FC for the autolysis proteome, while 2.42% at BFC vs. 1.31% at FC and 2.96 at BFC vs. 1.98 mol% at FC for the apoptosis proteome.

To achieve a detailed conclusion, each process has been treated separately from now on.

### 1. Autophagy

6 and 11 proteins out of the 95 autophagy proteins in *S. cerevisiae* were reported under BFC and FC, respectively. From the SGD GO Term Finder (Supplemental material 2), a high frequency of BFC autophagy proteome was found involved in reticulophagy —autophagy selective for the endoplasmic reticulum (2 proteins, Atg11p and Ypt1p; out of a total of 9 proteins annotated in the *S. cerevisiae* proteome). On the other hand, under FC, many proteins were associated to macroautophagy (also highly frequent if considering the total *S. cerevisiae* autophagy proteome, see Supplemental material 2) and organelle organization. This last biological process is referred to as the assembly, arrangement of constituent parts, or disassembly of an organelle within a cell, which are all frequent in growing cells and is the case of flor yeast under a nutrient-rich condition such as FC.

Yeast autophagy involves several steps: i) regulation of induction, ii) vesicle nucleation, iii) cargo packaging, iv) vesicle expansion and completion, v) retrieval, vi) docking and fusion, vii) vesicle breakdown and viii) permease efflux (Table 1 and Fig. 1). Two proteins (out of 26) with autophagy regulation function were quantified under BFC: Bcy1p and Pph21p (Fig. 2). The first inhibits protein kinase A (PKA) in the absence of cAMP (low levels when low glucose content) [36], that controls a variety of cellular processes and inhibits autophagy [3,37,38] while Pph21p, as well detected under FC, is a member of the Phosphatase 2A complex (PP2A) which is induced by TORC1 (one of the main regulators for autophagy along with PKA and Sch9 protein) and has an inhibitor function over the autophagosome formation genes. TORC1, PKA and Sch9 protein act as inhibitors of autophagosome formation. When autophagy triggering stimuli are perceived, these regulators are negatively induced [3,37,38].

**Figure 1.**
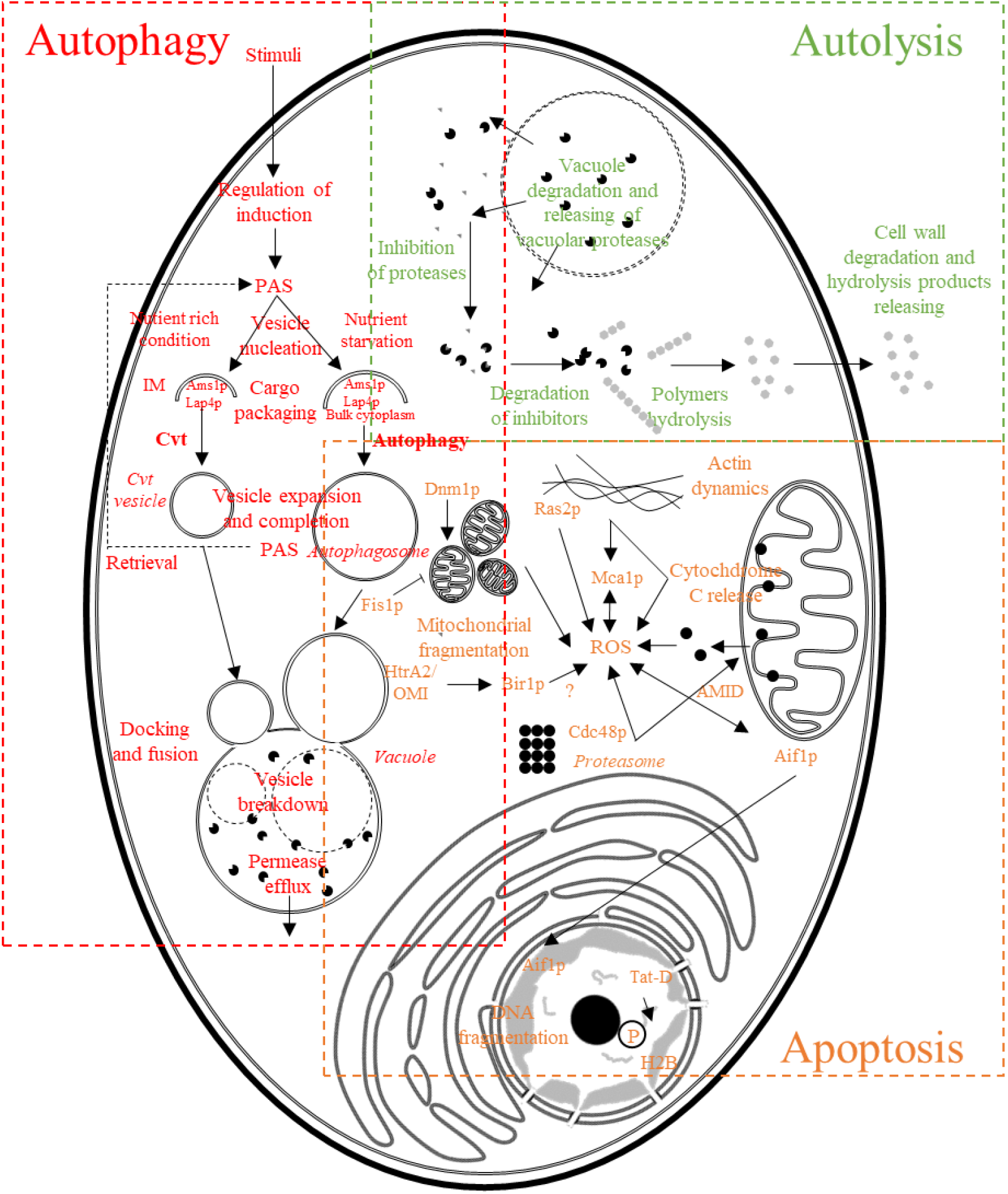
Illustration showing main steps in yeast autophagy, autolysis and apoptosis. PAS: pre-autophagosomal structure/phagophore assembly site; IM: isolation membrane for the formation of the sequestering vesicle; Cvt: cytoplasm to vacuole targeting; ROS: reactive oxygen species; AMID: AIF-homologous mitochondrion-associated inducer of death.

**Figure 2.**
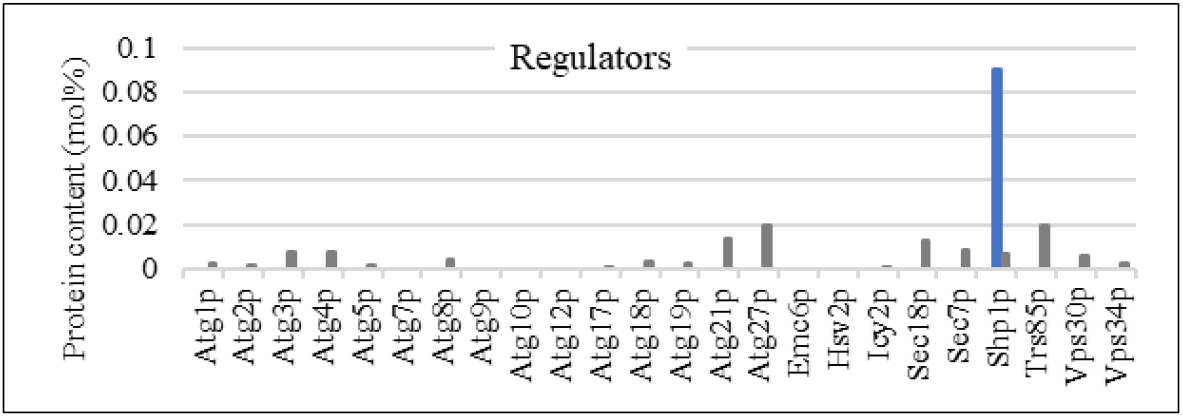
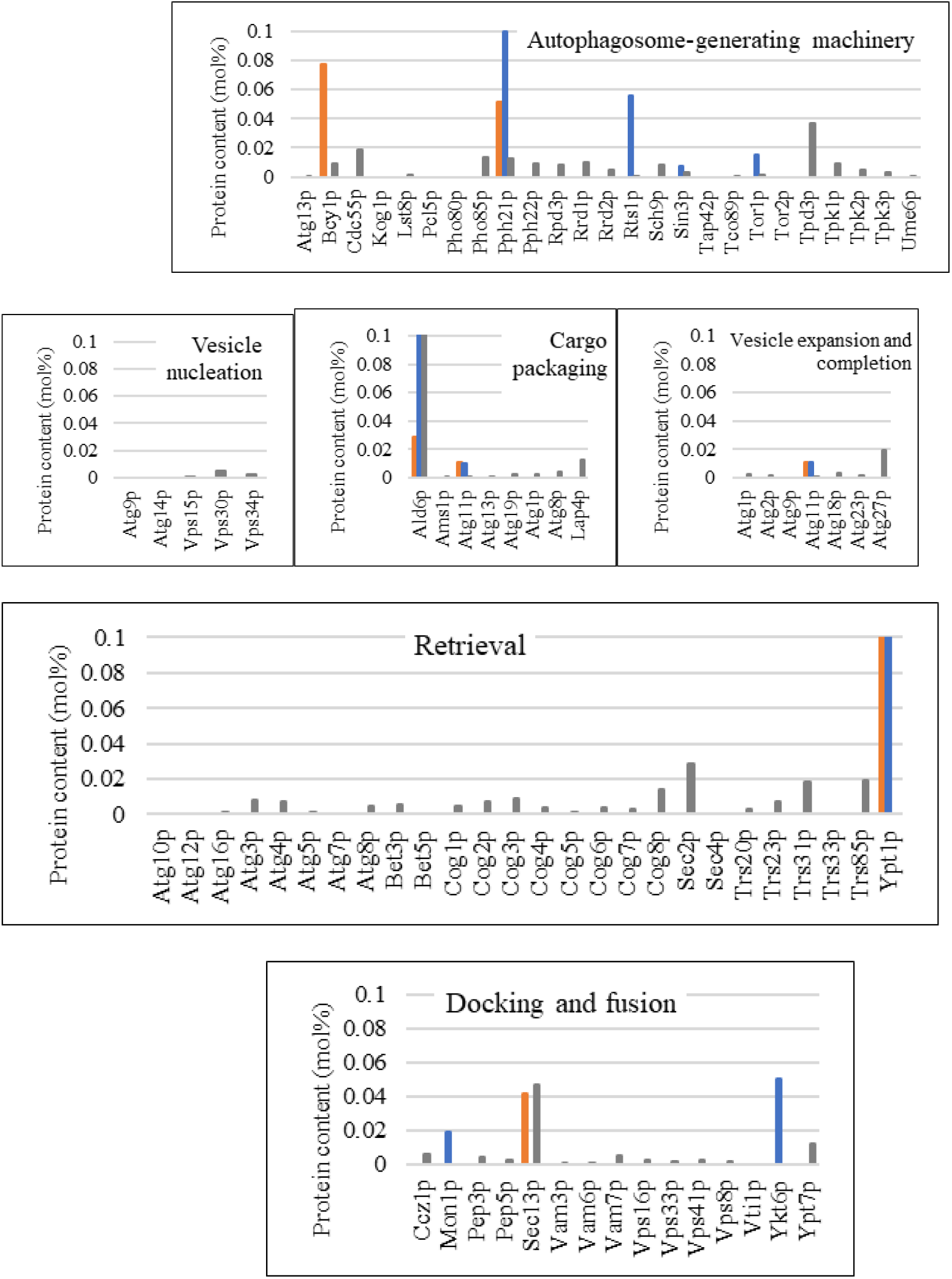
Content (mol%) of proteins related to autophagy detected in the flor yeast subjected to biofilm forming (BFC) in orange, fermentative conditions (FC) in blue and under nutrient rich conditions at log phase [31] in grey.

With regards to FC, besides Pph21p other proteins with autophagy regulation function were quantified: Rts1p, also member of the PP2A complex, Sin3p (also known as Rpd1p) component of both the Rpd3S and Rpd3L histone deacetylase complexes that regulate transcription, silencing, autophagy (as an inhibitor) and other processes by influencing chromatin remodeling [39–42]; and Tor1p, subunit of TORC1 (Fig. 2).

From the 24 proteins involved in the autophagosome formation process in the S. *cerevisiae* proteome, only 1 was detected under FC, Shp1p. In this step, the autophagosome-generating machinery comprised of Atg proteins collectively forms the pre-autophagosomal structure/phagophore assembly site (PAS) that will lead to the autophagosome vesicles. In addition, no proteins were found to take part in the vesicle nucleation step (initial stage of autophagosome vesicles formation) and two (out of 8 in *S. cerevisiae* proteome) in vesicle cargo packaging (Ald6p and Atg11p) under both BFC and FC. The cytosolic acetaldehyde dehydrogenase (Ald6p), specifically targeted to the vacuole by autophagosomes[22], was detected in high amounts under FC and log phase under rich conditions [31] (0.56 vs. 0.30 mol%, respectively) while under BFC remains much lower, 0.03 mol% (Fig. 2). The depletion of this protein has been used as a marker for the autophagy process [9,22]. Under nutrient starvation conditions, the Ald6p in cells was quickly depleted because of preferential degradation of this protein during autophagy.

Small GTPase Rab Ypt1p was the only protein quantified to be involved in vesicle expansion/completion out of the 26 reported in *S. cerevisiae*. Its content under BFC was 0.19 mol% while under FC, 0.32 mol%. This autophagy step is coordinately performed by Atg proteins (Atg3p-5p, Atg7p, Atg8p, Atg10p, Atg12p, Atg16p), Sec2/4p and Ypt1p and complexes COG and TRAPPIII (Fig. 2).

Atg11p, which participates in the cargo packaging step as well, was the unique protein reported in the study, having a role in the pre-autophagosomal structure retrieval (to form new autophagosomes). Atg11p together with Atg23p, and Atg27p, facilitates the anterograde transport of Atg9p to the PAS. This process occurs whether the cells are maintained in starvation state or growing state through the Cytoplasm-to-vacuole targeting (Cvt) pathway [43–45]. Cvt is a specific form and constitutive of autophagy that uses autophagosomal-like vesicles for selective transport of hydrolases Lap4p and Ams1p to the vacuole [46,47]. Depending on the nutrient condition, the vesicles engulf two different cargo: Ams1p and Lap4p (under nitrogen-rich conditions) and be besides these hydrolases, bulk cytoplasm (upon nutrient starvation) (shown in Fig. 1).

Out of the 15 proteins that mediate the docking and fusion of the autophagosome to the vacuole, one was reported under BFC, Sec13p and two, Mon1p and Ykt6p at FC (Fig. 2). This step results in the release of autophagic bodies that are further disintegrated, and their contents degraded for reuse in biosynthesis. Sec13p besides autophagy, is involved in other processes [48,49]. Meanwhile, Mon1p, in complex with Ccz1p (not identified), is required for multiple vacuole delivery pathways including the autophagy, pexophagy, endocytosis and cytoplasm-to-vacuole targeting (Cvt) pathway.

None of the proteins involved in vesicle breakdown and permease efflux have been detected in the present experiment.

Another gene found to be relevant in autophagy and whose product was observed in the present analysis under FC, is the AAA-type ATPase VPS4/CSC1 (Supplemental material 1). Vps4p is an AAA-type ATPase involved in multivesicular body protein sorting. Null mutant displays decreased autophagy while a gain-of-function mutant induces autophagy in rich medium [50,51].

### 2. Autolysis

Hydrolytic enzymes such as glucanases, proteases as well as nucleases play a major role in autolysis. Of all the enzymes involved, the activities of proteases have been the most extensively studied. According to Babayan et al. (1981) [52] yeast autolysis can be regarded as a four step process (Fig. 1): i) cell endostructures degradation and releasing vacuolar proteases in the cytoplasm, ii) inhibition of proteases and then activation due to the inhibitors degradation, iii) polymer hydrolysis and hydrolysis products accumulation in the cell, and iv) cell wall degradation and hydrolysis products releasing. Under both conditions, high frequencies of autolysis proteins were involved specifically in “protein catabolic process in the vacuole” GO Term (Supplemental material 2).

Among the 12 vacuolar proteases in *S. cerevisiae*, 4 were reported under BFC and 5 under FC; nevertheless, contents were much higher under BFC: 1.97 vs. 0.63 mol% being only 0.13 under nutrient-rich conditions (Table 1). Vacuolar proteases catalyze the non-specific degradation of cytoplasmic proteins, delocalized proteins from the secretory system, proteins delivered via autophagy, or plasma membrane proteins turned over via endocytosis [53,54]. Pep4p, the protein that most contributed to the mol% value in the BFC case, was quantified in 1.20 mol% which is over seven times higher than in FC, (0.16 mol%) (Fig. 3). Among the different types of proteases involved, Pep4p or Protease A is the main enzyme responsible for autolysis [55]. Lurton et al. (1989) [56] used specific proteases inhibitors to show that in acidic conditions, Pep4p was the principal enzyme involved in proteolysis during autolysis in a model wine system, despite numerous proteolytic enzymes present in yeast. This protein is required for posttranslational precursor maturation of other vacuolar proteinases, important for protein turnover after oxidative damage that may be occurring in BFC (flor yeast oxidative metabolism) and plays a protective role in acetic acid induced apoptosis [57–59]. Pep4p proteolytic activity is most efficient at acidic pH, as is the case of wines [60]. Some authors concluded that this protein is essential under conditions of nutrient starvation [58]. Alexandre et al. (2001) [55] support the idea that although protease A activity appeared to be responsible for peptides release, there is no clear correlation among protease A activity, cell death, and autolysis. It was suggested that protease A activity may be responsible for 80% of the nitrogen released during autolysis under optimum conditions. Using a Δpep4 mutant, Alexandre et al. (2001) [55] showed that protease A was responsible for 60% of the nitrogen released during autolysis in wine. These results suggest that other acidic proteases may also be involved in the proteolytic process. Consistent with this, Komano et al. (1999) and Olsen et al. (1999) [61,62] have identified other acidic proteases (Yapsin proteases Mkc7p, Yps1p, Yps3p, Yps6p and Yps7p) but none of them were reported in this proteomic approach. However, other proteins such as the vacuolar peptidases Ape3p (amino-) and Prc1p (carboxy-) were identified under BFC over the value quantified in FC, catalyzing the vacuole degradation that removes amino acids from the carboxy termini of non-specific proteins and small peptides [63].

**Figure 3.**
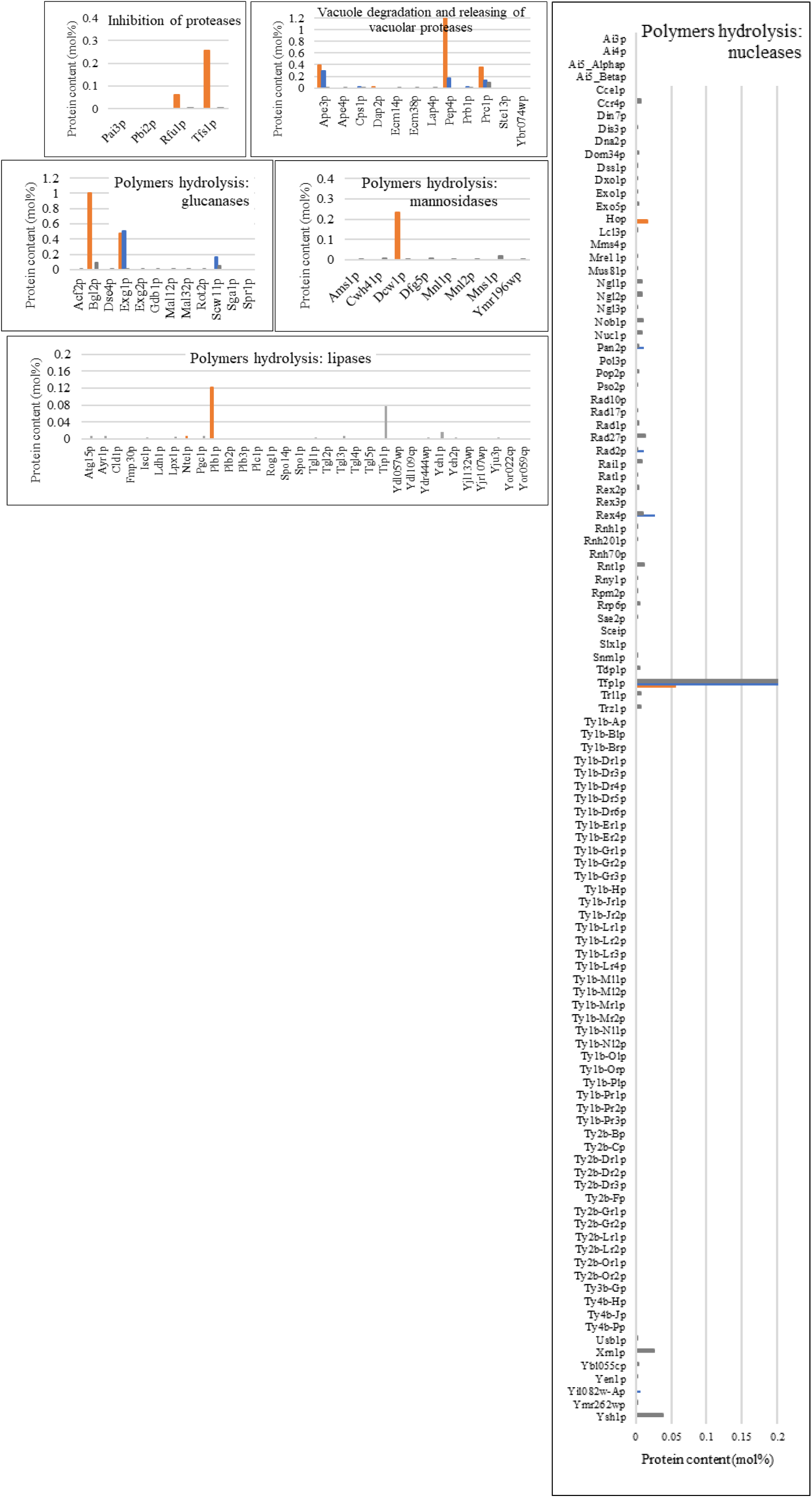
Content (mol%) of proteins related to autolysis detected in the flor yeast subjected to biofilm forming (BFC) in orange, fermentative conditions (FC) in blue and under nutrient rich conditions at log phase [31] in grey.

Alexandre et al. 2001 [55] showed that the proteolytic activity of yeast increases up to six-fold after sugar exhaustion, which is the case of BFC, but decreases when yeast cell autolysis starts. Also, temperature, pH and the yeast strain affect proteolytic activity during aging [1].

The released vacuolar proteases are initially inhibited by specific cytoplasmic inhibitors and are then activated due to their degradation. These inhibitors were only detected under BFC: Rfu1p with 0.06 mol% and Tfs1p with 0.26 mol% (Fig. 3). The first is the inhibitor of the Doa4p deubiquitinase (not reported) while Tfs1p is a specific and potent inhibitor of the vacuolar carboxypeptidase Y or Prc1p (quantified under BFC) [64,65]. During log phase growth, Tfs1p is found in the cytoplasm; it is re-localized to the vacuole in stationary phase [66,67]. Thus, as sampling was made at the middle of the log phase in both conditions, Tfs1p might be present in the cytoplasm exhibiting its inhibition function over the Prc1p released from the vacuole.

Besides proteases, glucanases and nucleases hydrolyze substrates under wine conditions [68,69]. In this approach, glucanases frequency was found higher under BFC (0.48 vs. 0.33%) but the difference was much bigger in terms of mol% content (1.48 vs. 0.67 mol%). Nucleases, on the other hand, showed the opposite trend: 0.48 vs. 0.82% at FC and BFC, respectively; and 0.07 and 0.36 mol% (Table 1). More FC nucleases (DNases and RNases) are explained since the yeasts have a higher cell division rate under FC where conditions are more favourable than under BFC for reproduction. As expected, under nutrient rich conditions at log phase where yeasts are not subjected to any stress, nucleases frequency and mol% values were higher than under BFC or FC [31].

Among BFC glucanases, cell wall enzyme endoglucanase Bgl2p reached a content of 1.01 mol% (not detected in FC) (Fig. 3), involved in beta-glucan degradation and also function biosynthetically as a transglycosylase [69]. It catalyzes the successive hydrolysis of beta-D-glucose units from the non-reducing ends of (1->3)-beta-D-glucans, releasing alpha-glucose. It is also involved in the incorporation of newly synthesized mannoprotein molecules into the cell wall and it introduces intrachain 1,6-beta linkages into 1,3-beta glucan, contributing to the rigid structure of the cell wall. Another glucanase quantified in high values under both conditions was the cell wall exoglucanase Exg1p (0.48 and 0.51 mol% in BFC and FC, respectively). This enzyme hydrolyzes both 1,3-beta- and 1,6-beta-linkages and even has beta-glucosidase activity. It could also function biosynthetically as a transglycosylase. This enzyme releases alpha-glucose. The endo-1,3-beta-D-glucosidase Scw11p was only identified under FC which is involved in cell separation and may play a role in conjugation during mating based on its regulation by Ste12p (not detected) [70].

It should be mentioned that hydrolytic products start to be released when their molecular masses are low enough to cross pores in the cell wall and that the cell wall degradation is not a requirement. During autolysis, the yeast cell wall degrades. Charpentier and Freyssinet (1989) [1] showed that cell wall degradation could be summarized as follows: first, glucans are hydrolysed by glucanases, thus releasing mannoproteins trapped or covalently linked to the glucans; second, the glucans are released due to either residual activities of cell wall glucanases or solubilized glucanases in the medium and finally, the protein fraction of the mannoproteins is degraded by proteolysis. Further, we looked for mannosidases in the proteome data set trying to find some differences among conditions. Only one was reported under BFC and none under FC. The one reported is Dcw1p that is localized in the cell membrane and may contribute to the mannose residues releasing from cell wall mannoproteins. Although proteases and glucanases degrade the cell wall, there is no breakdown of the cell wall [71]. The cell wall remains unbroken, with many ridges and folds, nevertheless the yeast cells have lost most of their cytoplasmic content.

With regards to the plasma membrane, its fate during this process is not clarified, however lipid release has been reported in sparkling wine aging [1]. In this study, only two lipases (more specifically lysophospholipases) have been quantified: Plb1p (0.12 mol%) under BFC and Nte1p under FC with only 0.01 mol%.

### 3. Apoptosis

Since the first description of apoptosis in yeasts [72], several yeast orthologues of crucial mammalian apoptotic proteins have been discovered [73–78], and conserved proteasomal, mitochondrial, and histone-regulated apoptotic pathways have been delineated (Fig. 1) [79–84]. Apoptosis involves three main steps: the perception of an external or internal signal, the signaling pathway phase and the execution phase that ends with the cell death. Apoptosis proteome in *S. cerevisiae* consists in 39 proteins.

Bir1p, Cpr3p, Kex1p, Mca1p, Pet9p, Por1p and Tdh2p were quantified under BFC 2-folding the FC content (Fig. 4). Por1p was the one showing the highest difference among both conditions: 0.28 vs. none, under BFC and FC, respectively. This is the mitochondrial outer membrane protein porin 1 which gene deletion (yeast voltage-dependent anion channel) enhances apoptosis triggered by acetic acid, H_2_O_2_ and diamide [84]. However, Liang and Zhou (2007) [85] proposed that this membrane protein enhances apoptosis in yeasts increasing resistance to apoptosis induced by Cu2+. Another protein showing significant differences in mol% that plays a central role in apoptosis, is Mca1p (see Fig. 4), reported under BFC with a content of 0.13 mol% while not identified under FC. It mediates apoptosis triggered by oxygen stress, salt stress or chronological aging or toxins and promotes the removal of insoluble protein aggregates during normal growth [73,86]. MCA1 plays a central role in yeast apoptosis, its deletion of the enhances the resistance against oxidative stress and delays age-induced cell death [73], although caspase-independent apoptosis occurs in yeast as well [87,88].

**Figure 4.**
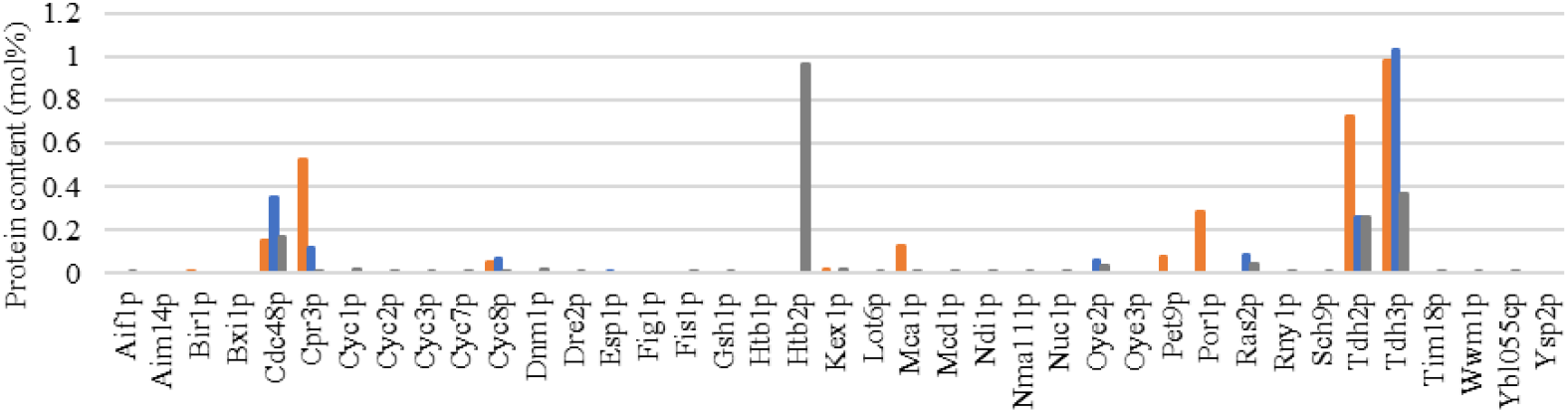
Content (mol%) of proteins related to apoptosis detected in the flor yeast subjected to biofilm forming (BFC) in orange, fermentative conditions (FC) in blue and under nutrient rich conditions at log phase [31] in grey.

Although with less content, another protein identified under BFC (0.08 mol%) and not under FC was Pet9p (Fig. 4). It catalyzes the exchange of ADP and ATP across the mitochondrial inner membrane. Genetic evidence indicates a possible role of the ADP/ATP carriers (AAC): Aac1p, Aac3p and Pet9p (Aac2p); in apoptosis [101]. Among them Pet9p is the major isoform of the translocator [89]. Pereira et al. (2007) [90] specifically pointed to a crucial role of AAC in yeast apoptosis as it is required for mitochondrial outer membrane permeabilization and cytochrome c release through the process (Fig. 1).

Two other proteins detected only under BFC but with less content were Bir1p and Kex1p. The first is an antiapoptotic protein that contains three Baculovirus IAP repeat domains, a protein motif which is usually found in inhibitor-of-apoptosis proteins [91] and appears to play independent roles in chromosome stability and apoptosis [77]. Kex1p, on the other hand, is a protease with a carboxypeptidase B-like function, involved in the C-terminal processing of the lysine and arginine residues from the precursors of K1, K2 and K28 killer toxins and a-factor (mating pheromone), in the programmed cell death caused by defective N-glycosylation and contributes to the active cell death program induced by acetic acid stress or during chronological aging [108].

Cpr3p, a yeast cyclophilin D homologue, was quantified under both conditions although in significantly higher content under BFC (0.52 vs. 0.12 mol%). Liang and Zhou (2007) [85] performed a genetic screen in which identified Cpr3p as activating the Cu2+ 371 -induced apoptotic program. Other protein folding the content under BFC is Tdh2p. Almeida et al. (2007)[4] by combining proteomic, genetic and biochemical approaches demonstrated that Nitric oxide (NO) and glyceraldehyde-3-phosphate dehydrogenase (GAPDH) as Tdh2p are crucial mediators of yeast H2O2-induced apoptosis, concluding that NO signaling and GAPDH S-nitrosation are linked with H2O2-induced apoptotic cell death. Evidence is presented showing that NO and GAPDH S-nitrosation also mediate cell death during chronological life span pointing to a physiological role of NO in yeast apoptosis. Further another GAPDH, Tdh3p was detected in very high amounts under both conditions (0.98 vs. 1.03 mol% under BFC and FC, respectively). The high presence of these proteins under FC could be explained since this protein is highly relevant in glycolysis which is essential under a typical fermentative condition.

Under FC, proteins like Oye2p and Ras2p were found specifically under this condition. The multifunctional protein Cdc48p doubled the content at this condition. Full length *OYE2* overexpression lowers endogenous reactive oxygen species (ROS), increases resistance to H2O2- induced programmed cell death (PCD) and significantly lowers ROS levels generated by organic prooxidants [92]. Reciprocally, oye2 yeast strains are sensitive to prooxidant-induced PCD. Odat O, et al. (2007) [92] firmly placed OYE proteins in the signaling network connecting ROS generation, PCD modulation and cytoskeletal dynamics in yeast (Fig. 1). Ras2p induces the production on ROS while Cdc48p is an antiapoptotic protein [93].

## 4. Discussion

The scarce proteins related to the autophagy process in both studied conditions (lower at BFC) along with the presence reported of the several autophagy inhibitors, points out a down- regulation of the autophagy genome in flor yeasts under BFC or FC. Further, the presence of Atg8p, an autophagy key protein that has been used as an experimental marker for autophagosomes, was neither quantified. The depletion of Ald6p is used as a marker for the autophagy process as it is specifically targeted to the vacuole by autophagosomes. Its lower amount in BFC compared with FC and log phase under rich-medium, represents an isolated fact that may indicate a progress in the autophagy process under BFC or, on the other hand, that the yeast stopped its synthesis at a certain point probably because its function is not relevant or is substituted by other Aldps such as Ald2p and Ald3p, which genes are both induced in response to ethanol or stress and repressed by glucose [94]. Porras-Agüera et al. (2020) [24] also reported a downregulation on the autophagy proteome in flor yeasts when subjected to sparkling wine-like conditions (i.e., CO2 overpressure and long aging time).

Flor yeasts under fermentative condition (FC) show higher values in frequency and content of autophagy proteins. Under a nutrient-rich condition such as FC, the autophagy role may be the reorganization of organelles, typical in growing cells, rather than material degradation that occurs in starving yeasts. The presence of Vps4p (relevant in autophagy) at this condition may indicate that autophagy is being induced at some extent. Piggot et al. (2011) [15] demonstrated that autophagy is induced early in wine fermentation in a nitrogen-replete environment, suggesting that autophagy may be triggered by other forms of stress that arise during fermentation. These authors also stated that autophagy genes are required for optimal survival throughout fermentation.

Autolysis and apoptosis proteome showed the opposite tendency of the autophagy in terms of frequency and protein content values, both higher under BFC. BFC vacuolar proteases triplicated those at FC in abundance while Pep4p, considered as the main responsible protein of the nitrogen release in wine autolysis [55], was the protein that most contributed to the content value in the autolysis BFC proteome, thus supporting other references that reported autolysis at biological aging [19]. Moreover, this protease may be active under BFC as the pH is acidic and there are no sugars [20,55]. Glucanases possibly play a role in cell expansion during growth, in cell-cell fusion during mating, and in spore release during sporulation. For this reason, these hydrolases might also be important under a condition with high growth rate, however, more glucanases in higher contents were reported under BFC pointing out that there is another process or are other processes that also requires this function (like autolysis). The high amounts of the cell wall glucanases Bgl2p and Exg1p in BFC can lead to cell wall glucans degradation. These two enzymatic families (vacuolar proteases and glucanases) were found to be significantly upregulated under sparkling wine-like conditions, wherein flor yeasts were subjected to prolonged nutrient starvation and CO2 overpressure [23].

Apoptosis proteins, as expected, were found more abundant under BFC than under FC, showing Cpr3p, Mca1p, Por1p, Tdh2/3p very high values. Under BFC, flor yeasts are subjected to a carbon starvation in which they are able to form a biofilm community. The self-destruction of damaged and old yeast cells, which consume dwindling nutrients, may contribute to the viability and reproductive success of healthier members of the community. The fact that high amounts of apoptosis activators, such as Mca1p or Cpr3p, were quantified, while none or very little amounts under FC, may point out that apoptosis is happening when the flor yeast is forming flor, which has never been reported before to our knowledge. Apoptotic death in yeast is suggested to be accompanied, at least under certain cases, by transfer of genetic material between cells [95]. This may be considered as a reason to explain why flor yeasts and fermentative yeasts differ genetically.

This study provides evidence about the autophagy, autolysis and apoptosis biological processes in flor yeasts when subjected to biofilm and fermentative conditions. However, besides proteomics, further works dealing with genetic approaches, deeper metabolomic analyses (including amino acids), transmission electron microscopy imaging, protein enzymatic activity and utilization of different flor yeast strains are required in order to achieve more solid conclusions. All said techniques could be considered and aimed to improve the knowledge of yeast behaviour under different enological conditions and further improve quality of wines.

Moreover, detecting apoptosis proteins in flor yeast biofilms highlights the potential use of these strains as unicellular eukaryotic models to study apoptosis for medical purposes.

## Supplementary Materials

The following supporting information can be downloaded at: www.mdpi.com/xxx/s1, Dataset S1: Proteins related to autophagy, autolysis and apoptosis detected in the flor yeast subjected to biofilm forming (BFC) and fermentative conditions (FC). This table includes: name of the protein, Uniprot accession number, name of the gene, brief description, biological process and molecular function, the molar weight (Mr), an identification score value (combination of the XCorr values for its constituent peptides), observable and observed peptides and relative content as calculated from its PAI value and mol/cell and mol% in *Saccharomyces cerevisiae* proteome measured under rich-medium conditions [31] as a reference. Dataset S2: Autophagy, autolysis and apoptosis proteins sorted in GO Terms of proteins related to autophagy, autolysis and apoptosis detected in the flor yeast subjected to biofilm forming (BFC) and fermentative conditions (FC). This table includes: GO term id and name, cluster frequency, background frequency, p-value, False Discovery Rate (FDR) and gene(s) annotated to the term. GO Terms in *Saccharomyces cerevisiae* were used as reference.

## Author Contributions

Conceptualization, J.M.G., J.C.M., and T.G.M.; methodology, J.M.G., J.C.M., and T.G.M.; software, J.M.G. and T.G.M.; validation, J.M.G., and J.C.M.; formal analysis, J.M.G.; investigation, J.M.G., J.C.M., and T.G.M.; resources, J.M.G., J.C.M., J.M., and T.G.M.; data curation, J.M.G., and J.C.M.; writing—original draft preparation, J.M.G.; writing—review and editing, J.C.M.; visualization, J.M.G., and J.C.M.; supervision, J.C.M., J.M., and T.G.M.; project administration, J.C.M., and J.M.; funding acquisition, J.C.M., J.M., and T.G.M. All authors have read and agreed to the published version of the manuscript.”

## Institutional Review Board Statement

Not applicable.

## Informed Consent Statement

Not applicable.

## Data Availability Statement

Data is contained within the article or supplementary material.

## Acknowledgments

The authors thank Minami Ogawa for her assistance with English language editing.

## Disclaimer/Publisher’s Note

The statements, opinions and data contained in all publications are solely those of the individual author(s) and contributor(s) and not of MDPI and/or the editor(s). MDPI and/or the editor(s) disclaim responsibility for any injury to people or property resulting from any ideas, methods, instructions or products referred to in the content.

## References

1. Alexandre, H.; Guilloux-Benatier, M. Yeast Autolysis in Sparkling Wine – a Review. Aust. J. Grape Wine Res. 2006, 12, 119–127, doi:10.1111/j.1755-0238.2006.tb00051.x.

2. Gnoinski, G.B.; Schmidt, S.A.; Close, D.C.; Goemann, K.; Pinfold, T.L.; Kerslake, F.L. Novel Methods to Manipulate Autolysis in Sparkling Wine: Effects on Yeast. Molecules 2021, 26, doi:10.3390/molecules26020387.

3. Budovskaya, Y.V.; Stephan, J.S.; Reggiori, F.; Klionsky, D.J.; Herman, P.K. The Ras/cAMP-Dependent Protein Kinase Signaling Pathway Regulates an Early Step of the Autophagy Process in Saccharomyces Cerevisiae. J. Biol. Chem. 2004, 279, 20663–20671, doi:10.1074/jbc.M400272200.

4. Almeida, B.; Buttner, S.; Ohlmeier, S.; Silva, A.; Mesquita, A.; Sampaio-Marques, B.; Osório, N.S.; Kollau, A.; Mayer, B.; Leão, C.;, et al. NO-Mediated Apoptosis in Yeast. J. Cell Sci. 2007, 120, 3279–3288, doi:10.1242/jcs.010926.

5. Reggiori, F.; Klionsky, D.J. Autophagic Processes in Yeast: Mechanism, Machinery and Regulation. Genetics 2013, 194, 341–361, doi:10.1534/genetics.112.149013.

6. Rego, A.; Trindade, D.; Chaves, S.R.; Manon, S.; Costa, V.; Sousa, M.J.; Côrte-Real, M. The Yeast Model System as a Tool towards the Understanding of Apoptosis Regulation by Sphingolipids. FEMS Yeast Res. 2014, 14, 160–178, doi:10.1111/1567-1364.12096.

7. Cebollero, E.; Carrascosa, A.V.; Gonzalez, R. Evidence for Yeast Autophagy during Simulation of Sparkling Wine Aging: A Reappraisal of the Mechanism of Yeast Autolysis in Wine. Biotechnol. Prog. 2005, 21, 614–616, doi:10.1021/bp049708y.

8. Babayan, T.L.; Bezrukov, M.G. Autolysis in yeasts. Acta Biotechnol. 1985, 5, 129–136, doi:10.1002/abio.370050205.

9. Cebollero, E.; Gonzalez, R. Induction of Autophagy by Second-Fermentation Yeasts during Elaboration of Sparkling Wines. Appl. Environ. Microbiol. 2006, 72, 4121–4127, doi:10.1128/AEM.02920-05.

10. Yousefi, S.; Simon, H.-U. Apoptosis Regulation by Autophagy Gene 5. Crit. Rev. Oncol. Hematol. 2007, 63, 241–244, doi:10.1016/j.critrevonc.2007.06.005.

11. Côrte-Real, M.; Madeo, F. Yeast Programed Cell Death and Aging. Front. Oncol. 2013, 3, 283, doi:10.3389/fonc.2013.00283.

12. Rangarajan, N.; Kapoor, I.; Li, S.; Drossopoulos, P.; White, K.K.; Madden, V.J.; Dohlman, H.G. Potassium Starvation Induces Autophagy in Yeast. J. Biol. Chem. 2020, 295, 14189–14202, doi:10.1074/jbc.RA120.014687.

13. Moreno-Arribas, V.; Pueyo, E.; Nieto, F.J.; Martín-Álvarez, P.J.; Polo, M.C. Influence of the Polysaccharides and the Nitrogen Compounds on Foaming Properties of Sparkling Wines. Food Chem. 2000, 70, 309–317, doi:10.1016/S0308-8146(00)00088-1.

14. Martinez-Rodriguez, A.J.; Gonzalez, R.; Carrascosa, A.V. Morphological Changes in Autolytic Wine Yeast during Aging in Two Model Systems. J. Food Sci. 2004, 69, M233*–* M239, doi:10.1111/j.1365-2621.2004.tb09893.x.

15. Piggott, N.; Cook, M.A.; Tyers, M.; Measday, V. Genome-Wide Fitness Profiles Reveal a Requirement for Autophagy During Yeast Fermentation. G3 2011, 1, 353–367, doi:10.1534/g3.111.000836.

16. Orozco, H.; Matallana, E.; Aranda, A. Genetic Manipulation of Longevity-Related Genes as a Tool to Regulate Yeast Life Span and Metabolite Production during Winemaking. Microb. Cell Fact. 2013, 12, 1, doi:10.1186/1475-2859-12-1.

17. Alexandre, H. Flor Yeasts of Saccharomyces Cerevisiae--Their Ecology, Genetics and Metabolism. Int. J. Food Microbiol. 2013, 167, 269–275, doi:10.1016/j.ijfoodmicro.2013.08.021.

18. Legras, J.-L.; Moreno-Garcia, J.; Zara, S.; Zara, G.; Garcia-Martinez, T.; Mauricio, J.C.; Mannazzu, I.; Coi, A.L.; Bou Zeidan, M.; Dequin, S.;, et al. Flor Yeast: New Perspectives Beyond Wine Aging. Front. Microbiol. 2016, 7, 503, doi:10.3389/fmicb.2016.00503.

19. Charpentier, C.; Dos Santos, A.M.; Feuillat, M. Release of Macromolecules by Saccharomyces Cerevisiae during Ageing of French Flor Sherry Wine “Vin Jaune.” Int. J. Food Microbiol. 2004, 96, 253–262, doi:10.1016/j.ijfoodmicro.2004.03.019.

20. Moreno-García, J.; García-Martínez, T.; Moreno, J.; Millán, M.C.; Mauricio, J.C. A Proteomic and Metabolomic Approach for Understanding the Role of the Flor Yeast Mitochondria in the Velum Formation. Int. J. Food Microbiol. 2014, 172, 21–29, doi:10.1016/j.ijfoodmicro.2013.11.030.

21. Xie, Z.; Nair, U.; Klionsky, D.J. Atg8 Controls Phagophore Expansion during Autophagosome Formation. Mol. Biol. Cell 2008, 19, 3290–3298, doi:10.1091/mbc.e07-12-1292.

22. Onodera, J.; Ohsumi, Y. Ald6p Is a Preferred Target for Autophagy in Yeast, Saccharomyces Cerevisiae*. J. Biol. Chem. 2004, 279, 16071–16076, doi:10.1074/jbc.M312706200.

23. Porras-Agüera, J.A.; Moreno-García, J.; Mauricio, J.C.; Moreno, J.; García-Martínez, T. First Proteomic Approach to Identify Cell Death Biomarkers in Wine Yeasts during Sparkling Wine Production. Microorganisms 2019, 7, doi:10.3390/microorganisms7110542.

24. Porras-Agüera, J.A.; Moreno-García, J.; González-Jiménez, M.D.C.; Mauricio, J.C.; Moreno, J.; García-Martínez, T. Autophagic Proteome in Two Saccharomyces Cerevisiae Strains During Second Fermentation for Sparkling Wine Elaboration. Microorganisms 2020, 8, doi:10.3390/microorganisms8040523.

25. Moreno-García, J.; García-Martínez, T.; Moreno, J.; Mauricio, J.C. Proteins Involved in Flor Yeast Carbon Metabolism under Biofilm Formation Conditions. Food Microbiol. 2015, 46, 25–33, doi:10.1016/j.fm.2014.07.001.

26. Moreno-García, J.; García-Martínez, T.; Millán, M.C.; Mauricio, J.C.; Moreno, J. Proteins Involved in Wine Aroma Compounds Metabolism by a Saccharomyces Cerevisiae Flor-Velum Yeast Strain Grown in Two Conditions. Food Microbiol. 2015, 51, 1–9, doi:10.1016/j.fm.2015.04.005.

27. Moreno-García, J.; Mauricio, J.C.; Moreno, J.; García-Martínez, T. Stress Responsive Proteins of a Flor Yeast Strain during the Early Stages of Biofilm Formation. Process Biochem. 2016, 51, 578–588, doi:10.1016/j.procbio.2016.02.011.

28. Moreno-García, J.; Mauricio, J.C.; Moreno, J.; García-Martínez, T. Differential Proteome Analysis of a Flor Yeast Strain under Biofilm Formation. Int. J. Mol. Sci. 2017, 18, doi:10.3390/ijms18040720.

29. Moreno-García, J.; Ogawa, M.; Joseph, C.M.L.; Mauricio, J.C.; Moreno, J.; García-Martínez, T. Comparative Analysis of Intracellular Metabolites, Proteins and Their Molecular Functions in a Flor Yeast Strain under Two Enological Conditions. World J. Microbiol. Biotechnol. 2018, 35, 6, doi:10.1007/s11274-018-2578-5.

30. Mauricio, J.C.; Moreno, J.J.; Ortega, J.M. In Vitro Specific Activities of Alcohol and Aldehyde Dehydrogenases from Two Flor Yeasts during Controlled Wine Aging. J. Agric. Food Chem. 1997, 45, 1967–1971, doi:10.1021/jf960634i.

31. Ghaemmaghami, S.; Huh, W.-K.; Bower, K.; Howson, R.W.; Belle, A.; Dephoure, N.; O’Shea, E.K.; Weissman, J.S. Global Analysis of Protein Expression in Yeast. Nature 2003, 425, 737–741, doi:10.1038/nature02046.

32. Salvadó, Z.; Chiva, R.; Rodríguez-Vargas, S.; Rández-Gil, F.; Mas, A.; Guillamón, J.M. Proteomic Evolution of a Wine Yeast during the First Hours of Fermentation. FEMS Yeast Res. 2008, 8, 1137–1146, doi:10.1111/j.1567-1364.2008.00389.x.

33. Gutiérrez, P.; Roldán, A.; Caro, I.; Pérez, L. Kinetic Study of the Velum Formation by Saccharomyces Cerevisiae (beticus Ssp.) during the Biological Aging of Wines. Process Biochem. 2010, 45, 493–499, doi:10.1016/j.procbio.2009.11.005.

34. Rodríguez, M.E.; Infante, J.J.; Mesa, J.J.; Rebordinos, L.; Cantoral, J.M. Enological Behaviour of Biofilms Formed by Genetically-Characterized Strains of Sherry Yeast. Open Biotechnol. J. 2013, 7, 23–29, doi:10.2174/1874070701307010023.

35. Ishihama, Y.; Oda, Y.; Tabata, T.; Sato, T.; Nagasu, T.; Rappsilber, J.; Mann, M. Exponentially Modified Protein Abundance Index (emPAI) for Estimation of Absolute Protein Amount in Proteomics by the Number of Sequenced Peptides per Protein. Mol. Cell. Proteomics 2005, 4, 1265–1272, doi:10.1074/mcp.M500061-MCP200.

36. Uno, I.; Matsumoto, K.; Ishikawa, T. Characterization of Cyclic AMP-Requiring Yeast Mutants Altered in the Regulatory Subunit of Protein Kinase. J. Biol. Chem. 1982, 257, 14110–14115, doi:10.1016/S0021-9258(19)45350-7.

37. Yorimitsu, T.; Klionsky, D.J. Endoplasmic Reticulum Stress: A New Pathway to Induce Autophagy. Autophagy 2007, 3, 160–162, doi:10.4161/auto.3653.

38. Noda, T.; Ohsumi, Y. Tor, a Phosphatidylinositol Kinase Homologue, Controls Autophagy in Yeast*. J. Biol. Chem. 1998, 273, 3963–3966, doi:10.1074/jbc.273.7.3963.

39. Silverstein, R.A.; Ekwall, K. Sin3: A Flexible Regulator of Global Gene Expression and Genome Stability. Curr. Genet. 2005, 47, 1–17, doi:10.1007/s00294-004-0541-5.

40. Carrozza, M.J.; Florens, L.; Swanson, S.K.; Shia, W.-J.; Anderson, S.; Yates, J.; Washburn, M.P.; Workman, J.L. Stable Incorporation of Sequence Specific Repressors Ash1 and Ume6 into the Rpd3L Complex. Biochim. Biophys. Acta 2005, 1731, 77–87; discussio*n* 75–76, doi:10.1016/j.bbaexp.2005.09.005.

41. Bernstein, B.E.; Tong, J.K.; Schreiber, S.L. Genomewide Studies of Histone Deacetylase Function in Yeast. Proc. Natl. Acad. Sci. U. S. A. 2000, 97, 13708–13713, doi:10.1073/pnas.250477697.

42. Aihara, M.; Jin, X.; Kurihara, Y.; Yoshida, Y.; Matsushima, Y.; Oku, M.; Hirota, Y.; Saigusa, T.; Aoki, Y.; Uchiumi, T.;, et al. Tor and the Sin3-Rpd3 Complex Regulate Expression of the Mitophagy Receptor Protein Atg32 in Yeast. J. Cell Sci. 2014, 127, 3184–3196, doi:10.1242/jcs.153254.

43. Shintani, T.; Klionsky, D.J. Cargo Proteins Facilitate the Formation of Transport Vesicles in the Cytoplasm to Vacuole Targeting Pathway. J. Biol. Chem. 2004, 279, 29889*–* 29894, doi:10.1074/jbc.M404399200.

44. He, C.; Song, H.; Yorimitsu, T.; Monastyrska, I.; Yen, W.-L.; Legakis, J.E.; Klionsky, D.J. Recruitment of Atg9 to the Preautophagosomal Structure by Atg11 Is Essential for Selective Autophagy in Budding Yeast. J. Cell Biol. 2006, 175, 925–935, doi:10.1083/jcb.200606084.

45. Cheong, H.; Nair, U.; Geng, J.; Klionsky, D.J. The Atg1 Kinase Complex Is Involved in the Regulation of Protein Recruitment to Initiate Sequestering Vesicle Formation for Nonspecific Autophagy in Saccharomyces Cerevisiae. Mol. Biol. Cell 2008, 19, 668–681, doi:10.1091/mbc.e07-08-0826.

46. Harding, T.M.; Hefner-Gravink, A.; Thumm, M.; Klionsky, D.J. Genetic and Phenotypic Overlap between Autophagy and the Cytoplasm to Vacuole Protein Targeting Pathway*. J. Biol. Chem. 1996, 271, 17621–17624, doi:10.1074/jbc.271.30.17621.

47. Yorimitsu, T.; Klionsky, D.J. Atg11 Links Cargo to the Vesicle-Forming Machinery in the Cytoplasm to Vacuole Targeting Pathway. Mol. Biol. Cell 2005, 16, 1593–1605, doi:10.1091/mbc.e04-11-1035.

48. Bonifacino, J.S.; Glick, B.S. The Mechanisms of Vesicle Budding and Fusion. Cell 2004, 116, 153–166, doi:10.1016/s0092-8674(03)01079-1.

49. Dokudovskaya, S.; Waharte, F.; Schlessinger, A.; Pieper, U.; Devos, D.P.; Cristea, I.M.; Williams, R.; Salamero, J.; Chait, B.T.; Sali, A.;, et al. A Conserved Coatomer-Related Complex Containing Sec13 and Seh1 Dynamically Associates with the Vacuole in Saccharomyces Cerevisiae. Mol. Cell. Proteomics 2011, 10, M110.006478, doi:10.1074/mcp.M110.006478.

50. Shirahama, K.; Noda, T.; Ohsumi, Y. Mutational Analysis of Csc1/Vps4p: Involvement of Endosome in Regulation of Autophagy in Yeast. Cell Struct. Funct. 1997, 22, 501–509, doi:10.1247/csf.22.501.

51. Nebauer, R.; Rosenberger, S.; Daum, G. Phosphatidylethanolamine, a Limiting Factor of Autophagy in Yeast Strains Bearing a Defect in the Carboxypeptidase Y Pathway of Vacuolar Targeting. J. Biol. Chem. 2007, 282, 16736–16743, doi:10.1074/jbc.M611345200.

52. Babayan, T.L.; Bezrukov, M.G.; Latov, V.K.; Belikov, V.M.; Belavtseva, E.M.; Titova, E.F. Induced Autolysis ofSaccharomyces Cerevisiae: Morphological Effects, Rheological Effects, and Dynamics of Accumulation of Extracellular Hydrolysis Products. Curr. Microbiol. 1981, 5, 163–168, doi:10.1007/bf01578522.

53. Chiang, H.L.; Schekman, R. Regulated Import and Degradation of a Cytosolic Protein in the Yeast Vacuole. Nature 1991, 350, 313–318, doi:10.1038/350313a0.

54. Klionsky, D.J.; Emr, S.D. Autophagy as a Regulated Pathway of Cellular Degradation. Science 2000, 290, 1717–1721, doi:10.1126/science.290.5497.1717.

55. Alexandre, H.; Heintz, D.; Chassagne, D.; Guilloux-Benatier, M.; Charpentier, C.; Feuillat, M. Protease A Activity and Nitrogen Fractions Released during Alcoholic Fermentation and Autolysis in Enological Conditions. J. Ind. Microbiol. Biotechnol. 2001, 26, 235–240, doi:10.1038/sj.jim.7000119.

56. Lurton, L.; Segain, J.P.; Feuillat, M. Etude de La Protéolyse Au Cours de L’autolyse de Levures Au Milieu Acide. Sci. Aliments 1989, 9, 111–123.

57. Wolff, A.M.; Din, N.; Petersen, J.G. Vacuolar and Extracellular Maturation of Saccharomyces Cerevisiae Proteinase A. Yeast 1996, 12, 823–832, doi:10.1002/(SICI)1097-0061(199607)12:9<823::AID-YEA975>3.0.CO;2-J.

58. Marques, M.; Mojzita, D.; Amorim, M.A.; Almeida, T.; Hohmann, S.; Moradas-Ferreira, P.; Costa, V. The Pep4p Vacuolar Proteinase Contributes to the Turnover of Oxidized Proteins but PEP4 Overexpression Is Not Sufficient to Increase Chronological Lifespan in Saccharomyces Cerevisiae. Microbiology 2006, 152, 3595–3605, doi:10.1099/mic.0.29040-0.

59. Pereira, H.; Azevedo, F.; Rego, A.; Sousa, M.J.; Chaves, S.R.; Côrte-Real, M. The Protective Role of Yeast Cathepsin D in Acetic Acid-Induced Apoptosis Depends on ANT (Aac2p) but Not on the Voltage-Dependent Channel (Por1p). FEBS Lett. 2013, 587, 200–205, doi:10.1016/j.febslet.2012.11.025.

60. Sørensen, S.O.; van den Hazel, H.B.; Kielland-Brandt, M.C.; Winther, J.R. pH-Dependent Processing of Yeast Procarboxypeptidase Y by Proteinase A in Vivo and in Vitro. Eur. J. Biochem. 1994, 220, 19–27, doi:10.1111/j.1432-1033.1994.tb18594.x.

61. Komano, H.; Rockwell, N.; Wang, G.T.; Krafft, G.A.; Fuller, R.S. Purification and Characterization of the Yeast Glycosylphosphatidylinositol-Anchored, Monobasic-Specific Aspartyl Protease Yapsin 2 (Mkc7p)*. J. Biol. Chem. 1999, 274, 24431–24437, doi:10.1074/jbc.274.34.24431.

62. Olsen, V.; Cawley, N.X.; Brandt, J.; Egel-Mitani, M.; Loh, Y.P. Identification and Characterization of Saccharomyces Cerevisiae Yapsin 3, a New Member of the Yapsin Family of Aspartic Proteases Encoded by the YPS3 Gene. Biochem. J 1999, 339 *( Pt* *2*), 407–411.

63. Van Den Hazel, H.B.; Kielland-Brandt, M.C.; Winther, J.R. Review: Biosynthesis and Function of Yeast Vacuolar Proteases. Yeast 1996, 12, 1–16, doi:10.1002/(sici)1097-0061(199601)12:1<1::aid-yea902>3.0.co;2-n.

64. Papa, F.R.; Amerik, A.Y.; Hochstrasser, M. Interaction of the Doa4 Deubiquitinating Enzyme with the Yeast 26S Proteasome. Mol. Biol. Cell 1999, 10, 741–756, doi:10.1091/mbc.10.3.741.

65. Bruun, A.W.; Svendsen, I.; Sørensen, S.O.; Kielland-Brandt, M.C.; Winther, J.R. A High-Affinity Inhibitor of Yeast Carboxypeptidase Y Is Encoded by TFS1 and Shows Homology to a Family of Lipid Binding Proteins. Biochemistry 1998, 37, 3351–3357, doi:10.1021/bi971286w.

66. Mima, J.; Fukada, H.; Nagayama, M.; Ueda, M. Specific Membrane Binding of the Carboxypeptidase Y Inhibitor I(C), a Phosphatidylethanolamine-Binding Protein Family Member. FEBS J. 2006, 273, 5374–5383, doi:10.1111/j.1742-4658.2006.05530.x.

67. Fukada, H.; Mima, J.; Nagayama, M.; Kato, M.; Ueda, M. Biochemical Analysis of the Yeast Proteinase Inhibitor (IC) Homolog ICh and Its Comparison with IC. Biosci. Biotechnol. Biochem. 2007, 71, 472–480, doi:10.1271/bbb.60528.

68. Martínez-Rodríguez, A.J.; Carrascosa, A.V.; Martín-Álvarez, P.J.; Moreno-Arribas, V.; Polo, M.C. Influence of the Yeast Strain on the Changes of the Amino Acids, Peptides and Proteins during Sparkling Wine Production by the Traditional Method. J. Ind. Microbiol. Biotechnol. 2002, 29, 314–322, doi:10.1038/sj.jim.7000323.

69. Zhao, J.; Fleet, G.H. Degradation of RNA during the Autolysis of Saccharomyces Cerevisiae Produces Predominantly Ribonucleotides. J. Ind. Microbiol. Biotechnol. 2005, 32, 415–423, doi:10.1007/s10295-005-0008-9.

70. Zeitlinger, J.; Simon, I.; Harbison, C.T.; Hannett, N.M.; Volkert, T.L.; Fink, G.R.; Young, R.A. Program-Specific Distribution of a Transcription Factor Dependent on Partner Transcription Factor and MAPK Signaling. Cell 2003, 113, 395–404, doi:10.1016/s0092-8674(03)00301-5.

71. Vosti, D.C.; Joslyn, M.A. Autolysis of Baker’s Yeast. Appl. Microbiol. 1954, 2, 70–78, doi:10.1128/am.2.2.70-78.1954.

72. Madeo, F.; Fröhlich, E.; Fröhlich, K.U. A Yeast Mutant Showing Diagnostic Markers of Early and Late Apoptosis. J. Cell Biol. 1997, 139, 729–734, doi:10.1083/jcb.139.3.729.

73. Madeo, F.; Herker, E.; Maldener, C.; Wissing, S.; Lächelt, S.; Herlan, M.; Fehr, M.; Lauber, K.; Sigrist, S.J.; Wesselborg, S.;, et al. A Caspase-Related Protease Regulates Apoptosis in Yeast. Mol. Cell 2002, 9, 911–917, doi:10.1016/s1097-2765(02)00501-4.

74. Wissing, S.; Ludovico, P.; Herker, E.; Büttner, S.; Engelhardt, S.M.; Decker, T.; Link, A.; Proksch, A.; Rodrigues, F.; Corte-Real, M.;, et al. An AIF Orthologue Regulates Apoptosis in Yeast. J. Cell Biol. 2004, 166, 969–974, doi:10.1083/jcb.200404138.

75. Qiu, J.; Yoon, J.-H.; Shen, B. Search for Apoptotic Nucleases in Yeast: ROLE OF Tat- D NUCLEASE IN APOPTOTIC DNA DEGRADATION*. J. Biol. Chem. 2005, 280, 15370*–* 15379, doi:10.1074/jbc.M413547200.

76. Li, W.; Sun, L.; Liang, Q.; Wang, J.; Mo, W.; Zhou, B. Yeast AMID Homologue Ndi1p Displays Respiration-Restricted Apoptotic Activity and Is Involved in Chronological Aging. Mol. Biol. Cell 2006, 17, 1802–1811, doi:10.1091/mbc.e05-04-0333.

77. Walter, D.; Wissing, S.; Madeo, F.; Fahrenkrog, B. The Inhibitor-of-Apoptosis Protein Bir1p Protects against Apoptosis in S. Cerevisiae and Is a Substrate for the Yeast Homologue of Omi/HtrA2. J. Cell Sci. 2006, 119, 1843–1851, doi:10.1242/jcs.02902.

78. Fahrenkrog, B. Nma111p, the pro-Apoptotic HtrA-like Nuclear Serine Protease in Saccharomyces Cerevisiae: A Short Survey. Biochem. Soc. Trans. 2011, 39, 1499–1501, doi:10.1042/BST0391499.

79. Ligr, M.; Velten, I.; Fröhlich, E.; Madeo, F.; Ledig, M.; Fröhlich, K.U.; Wolf, D.H.; Hilt, W. The Proteasomal Substrate Stm1 Participates in Apoptosis-like Cell Death in Yeast. Mol. Biol. Cell 2001, 12, 2422–2432, doi:10.1091/mbc.12.8.2422.

80. Ludovico, P.; Rodrigues, F.; Almeida, A.; Silva, M.T.; Barrientos, A.; Côrte-Real, M. Cytochrome c Release and Mitochondria Involvement in Programmed Cell Death Induced by Acetic Acid in Saccharomyces Cerevisiae. Mol. Biol. Cell 2002, 13, 2598–2606, doi:10.1091/mbc.e01-12-0161.

81. Fannjiang, Y.; Cheng, W.-C.; Lee, S.J.; Qi, B.; Pevsner, J.; McCaffery, J.M.; Hill, R.B.; Basañez, G.; Hardwick, J.M. Mitochondrial Fission Proteins Regulate Programmed Cell Death in Yeast. Genes Dev. 2004, 18, 2785–2797, doi:10.1101/gad.1247904.

82. Ahn, S.-H.; Cheung, W.L.; Hsu, J.-Y.; Diaz, R.L.; Smith, M.M.; Allis, C.D. Sterile 20 Kinase Phosphorylates Histone H2B at Serine 10 during Hydrogen Peroxide-Induced Apoptosis in S. Cerevisiae. Cell 2005, 120, 25–36, doi:10.1016/j.cell.2004.11.016.

83. Gourlay, C.W.; Ayscough, K.R. Identification of an Upstream Regulatory Pathway Controlling Actin-Mediated Apoptosis in Yeast. J. Cell Sci. 2005, 118, 2119–2132, doi:10.1242/jcs.02337.

84. Pozniakovsky, A.I.; Knorre, D.A.; Markova, O.V.; Hyman, A.A.; Skulachev, V.P.; Severin, F.F. Role of Mitochondria in the Pheromone- and Amiodarone-Induced Programmed Death of Yeast. J. Cell Biol. 2005, 168, 257–269, doi:10.1083/jcb.200408145.

85. Liang, Q.; Zhou, B. Copper and Manganese Induce Yeast Apoptosis via Different Pathways. Mol. Biol. Cell 2007, 18, 4741–4749, doi:10.1091/mbc.e07-05-0431.

86. Lee, R.E.C.; Brunette, S.; Puente, L.G.; Megeney, L.A. Metacaspase Yca1 Is Required for Clearance of Insoluble Protein Aggregates. Proc. Natl. Acad. Sci. U. S. A. 2010, 107, 13348–13353, doi:10.1073/pnas.1006610107.

87. Guscetti, F.; Nath, N.; Denko, N. Functional Characterization of Human Proapoptotic Molecules in Yeast S. Cerevisiae. FASEB J. 2005, 19, 464–466, doi:10.1096/fj.04-2316fje.

88. Zhang, N.-N.; Dudgeon, D.D.; Paliwal, S.; Levchenko, A.; Grote, E.; Cunningham, K.W. Multiple Signaling Pathways Regulate Yeast Cell Death during the Response to Mating Pheromones. Mol. Biol. Cell 2006, 17, 3409–3422, doi:10.1091/mbc.e06-03-0177.

89. Smith, C.P.; Thorsness, P.E. The Molecular Basis for Relative Physiological Functionality of the ADP/ATP Carrier Isoforms in Saccharomyces Cerevisiae. Genetics 2008, 179, 1285–1299, doi:10.1534/genetics.108.087700.

90. Pereira, C.; Camougrand, N.; Manon, S.; Sousa, M.J.; Côrte-Real, M. ADP/ATP Carrier Is Required for Mitochondrial Outer Membrane Permeabilization and Cytochrome c Release in Yeast Apoptosis. Mol. Microbiol. 2007, 66, 571–582, doi:10.1111/j.1365-2958.2007.05926.x.

91. Uren, A.G.; Beilharz, T.; O’Connell, M.J.; Bugg, S.J.; van Driel, R.; Vaux, D.L.; Lithgow, T. Role for Yeast Inhibitor of Apoptosis (IAP)-like Proteins in Cell Division. Proc. Natl. Acad. Sci. U. S. A. 1999, 96, 10170–10175, doi:10.1073/pnas.96.18.10170.

92. Odat, O.; Matta, S.; Khalil, H.; Kampranis, S.C.; Pfau, R.; Tsichlis, P.N.; Makris, A.M. Old Yellow Enzymes, Highly Homologous FMN Oxidoreductases with Modulating Roles in Oxidative Stress and Programmed Cell Death in Yeast*. J. Biol. Chem. 2007, 282, 36010*–* 36023, doi:10.1074/jbc.M704058200.

93. Büttner, S.; Eisenberg, T.; Herker, E.; Carmona-Gutierrez, D.; Kroemer, G.; Madeo, F. Why Yeast Cells Can Undergo Apoptosis: Death in Times of Peace, Love, and War. J. Cell Biol. 2006, 175, 521–525, doi:10.1083/jcb.200608098.

94. Wang, X.; Mann, C.J.; Bai, Y.; Ni, L.; Weiner, H. Molecular Cloning, Characterization, and Potential Roles of Cytosolic and Mitochondrial Aldehyde Dehydrogenases in Ethanol Metabolism in Saccharomyces Cerevisiae. J. Bacteriol. 1998, 180, 822–830, doi:10.1128/JB.180.4.822-830.1998.

95. Diker-Cohen, T.; Koren, R.; Ravid, A. Programmed Cell Death of Stressed Keratinocytes and Its Inhibition by Vitamin D: The Role of Death and Survival Signaling Pathways. Apoptosis 2006, 11, 519–534, doi:10.1007/s10495-006-5115-1.

